# HIWI2 Influences Endosomal Trafficking and Eph Receptor Signaling in Photoreceptor Cells

**DOI:** 10.64898/2026.03.09.710476

**Authors:** Rupa Roy, Rahini Rajendran, Jayamuruga Pandian Arunachalam, Subbulakshmi Chidambaram

## Abstract

Photoreceptor integrity depends on the precise coordination of membrane trafficking and signal transmission. Despite their well-known roles in germline biology, the functions of PIWI family proteins in post-mitotic neuronal cells remain unclear. We investigated the role of HIWI2 in photoreceptor-derived 661W cells. Silencing of HIWI2 resulted in a significant decrease in the early endosomal marker, Rab5, and its effector EEA1, and reduced expression of the recycling endosome marker Rab11, indicating poor endosomal sorting and receptor recycling. In contrast, the marker for late endosomes, Rab7, was significantly upregulated, suggesting a shift toward degradative trafficking pathways, in line with increased receptor breakdown. These trafficking shifts led to the degradation of EphA2 and EphB2 receptors, as confirmed by a phospho-proteome receptor tyrosine kinase array and further supported by immunoblotting, and were accompanied by a compensatory increase in Akt phosphorylation. Furthermore, HIWI2 deficiency impaired cell motility in wound-healing assays. These results propose HIWI2 as a critical regulator of endosomal sorting and Eph receptor stability, providing a novel link between the PIWI pathway and photoreceptor integrity.

## Introduction

Photoreceptor (PR) cells are highly specialized, photosensitive neurons that transduce light into an electrical signal which are then relayed to the brain through bipolar cells and retinal ganglion cells. PR outer segments are in contact with the retinal pigmented epithelium (RPE) through a specialized extracellular matrix (ECM), the inter-photoreceptor matrix (IPM) [1]. PRs are highly energetic cells whose survival relies on proper supply of nutrients and energy production [2–4]. This is achieved by various proteins and receptor tyrosine kinases (RTKs). PR consists of specialized proteins involved in phototransduction, including opsin, transducin, rhodopsin kinase, arrestin, and specialized RTKs [5,6]. PR degeneration is a major cause of irreversible blindness, characterized by progressive structural and functional deterioration of PR cells. The highly polarized architecture of PR, along with the renewal of their outer segments, relies on intensive vesicular transport, continuous membrane turnover, and signaling through receptors. An impairment in the performance of these processes disrupts cellular homeostasis and directly contributes to retinal degeneration [7].

RTKs are a subclass of tyrosine kinases that are involved in most aspects of cell fate determination, differentiation, patterning, proliferation, growth, and survival [8,9]. The various functions of RTKs are programmed to take place at precisely controlled locations and times both during development and in adult tissue homeostasis. Indeed, the spatiotemporal regulation of receptor abundance, as well as the availability and density at the membrane are all major determinants of RTK output [10,11]. A crucial RTK, Ephs belongs to the largest RTK-subfamily and are involved in retinal patterning, cell positioning, synaptic organization, and cytoskeletal remodeling [12,13]. Loss of Ephs is associated with defective projection of photoreceptor axons as well as aberrant targeting and loss of medulla and lobular cortical axons [14,15]. All these studies imply that RTK signaling is significant. The trafficking of RTKs is tightly regulated by Rab GTPases. Rab5 ensures ligand-augmented activation, Rab11 facilitates receptor recycling, and Rab7 mediates degradation [16–17]. Thus, disturbances in Rab homeostasis can profoundly alter RTK signaling, including Eph-mediated pathways.

Rab GTPases regulates vesicular trafficking by modulating the transport of cargo between the early, recycling, and late endosomal compartments [18]. More precisely, Rab5 and its effector EEA1 direct membrane fusion and sorting of cargo into the early endosomes. Recycling endosomes that function to reintegrate membrane proteins, such as components of phototransduction and surface receptors, are modulated by Rab11 [19,20]. Rab7 controls trafficking into the late endosome-lysosome pathway with an additional role in degradation [21]. Dysregulation of each of these Rab-mediated pathways has been directly implicated in photoreceptor degeneration: perturbations of Rab5 result in the accumulation and mislocalization of opsins on endosomes [22]; loss of Rab11 leads to defective outer segment maintenance [19]; and dysregulation of Rab7 contributes to accumulation of rhodopsin and results in late-stage degeneration [23].

RNA-binding proteins (RBPs) are now recognized as major players in the regulation of trafficking in neurons, the mobility of receptors, and the cytoskeleton dynamics [24–27]. Among these, the PIWI proteins such as HIWI2 are known to pair with piRNAs and are involved in the stability of RNA, translation, and stress responses [28–30]. HIWI2 is expressed in retinal pigment epithelium (RPE) and is altered in oxidative stress and diabetic retinopathy [31–34]. However, its functional role in PR remains elusive.

Here, we identify HIWI2 as a critical influencer of photoreceptor vesicular trafficking and RTK/Eph signaling. We demonstrated that HIWI2 depletion disrupts Rab expression, shifts endosomal balance toward degradation, broadly suppresses RTK activation, and specifically impairs EphA2 and EphB2 signaling. This underlying molecular alteration led to Akt hyperactivation. Our findings reveal a previously unrecognized regulatory axis essential for PR function.

## Methods

### Cell culture

661W cells (Photoreceptor cells) were a kind gift given by Dr. Muayyad Al-Ubaidi (Houston University) [35,36]. 661W-cells were cultured using DMEM-F12 media (Sigma-Aldrich) supplemented with 10% (*v*/v) fetal bovine serum (FBS; Gibco). The cells were maintained in 5% CO_2_ at 37 °C with antibiotics and antimycotics (HIMEDIA).

### RNAi experiments

The 661W cells were grown to confluency and 2X10^5^ cells were seeded in 6-well plates. DsiRNA specific for HIWI2 (sense strand 5′-GCAUCACUAGAUGGACAAUCCAAGA-3′; antisense3’-ACCGUAGUGAUCUACCUGUUAGGUUCU-5′; Integrated DNA Technologies) was transfected to the cells using X-tremeGENE siRNA Transfection (Sigma-Aldrich) as per the manufacturer’s protocol. Protein lysates were obtained by lysing the cells 48 hours post-transfection.

### Western blot

Cells were lysed using radioimmunoprecipitation assay buffer (RIPA) with protease inhibitor cocktail (Roch). 35μg of protein was resolved on SDS-PAGE gel and electro-transferred to nitrocellulose membrane (GE Healthcare). The blots were incubated in blocking buffer (5% BSA) for 1 h and were probed against HIWI2 (Abcam), EphrinA2-total (CST), EphrinA2-phospho (CST), EphrinB2-total (CST), EphrinB2-phospho (Thermo Scientific), Rab5 (CST), Rab11 (CST), Early Endosome antigen-1 (CST), Rab 7 (CST), Actin (Sigma), GAPDH (CST), primary antibodies in a 1:1000 dilution. Anti-rabbit (CST) and anti-mouse secondary antibodies (CST) were used in a 1:10,000 dilution. The blots were developed in Fusion Solo S (Vilber-Germany) using an enhanced chemiluminescence reagent (Pierce Western Blotting Substrate; ThermoScientific).

### Proteome profiler array

Proteins that were altered after HIWI2 silencing were screened using the Phospho-RTK array kit (ARY001B, CST). Protein lysates (200μg) were prepared according to manufacturer’s protocol and was used for the array. The array was imaged in Fusion Solo S (Vilber-Germany).

### Statistical analysis

Student’s *t*-test was used to comparatively analyze the mean obtained from three independent experiments. The difference between the samples was considered to be statistically significant when the *p*-values were *p* < 0.05.

## Results

### 1. Suppression of HIWI2 leads to altered expression of endosomal Rabs

To investigate whether HIWI2 regulates vesicular trafficking, we first analyzed Rab proteins involved in endosomal progression. Silencing of HIWI2 resulted in significant reductions in Rab5 (2-fold) and its early endosome effector, EEA1 (6-fold) (Fig.1A-C) indicating compromised early endosomal maturation. We next assessed Rab11, a key recycling endosome marker. HIWI2 depletion led to a profound reduction in Rab11 expression (8.6-fold) (Fig.1D), suggesting defective membrane protein recycling. In contrast to Rab5 and Rab11, the late endosomal marker Rab7 was increased by 1.8-fold in HIWI2-silenced cells (Fig.1E), indicating an accelerated step from late endosomes to lysosomes. The complementary expression of reduced early/recycling endosomal markers and increased Rab7 levels suggests that HIWI2 maintains endosomal integrity and that its downregulation promotes a bias towards degradation.

**Fig. 1.**
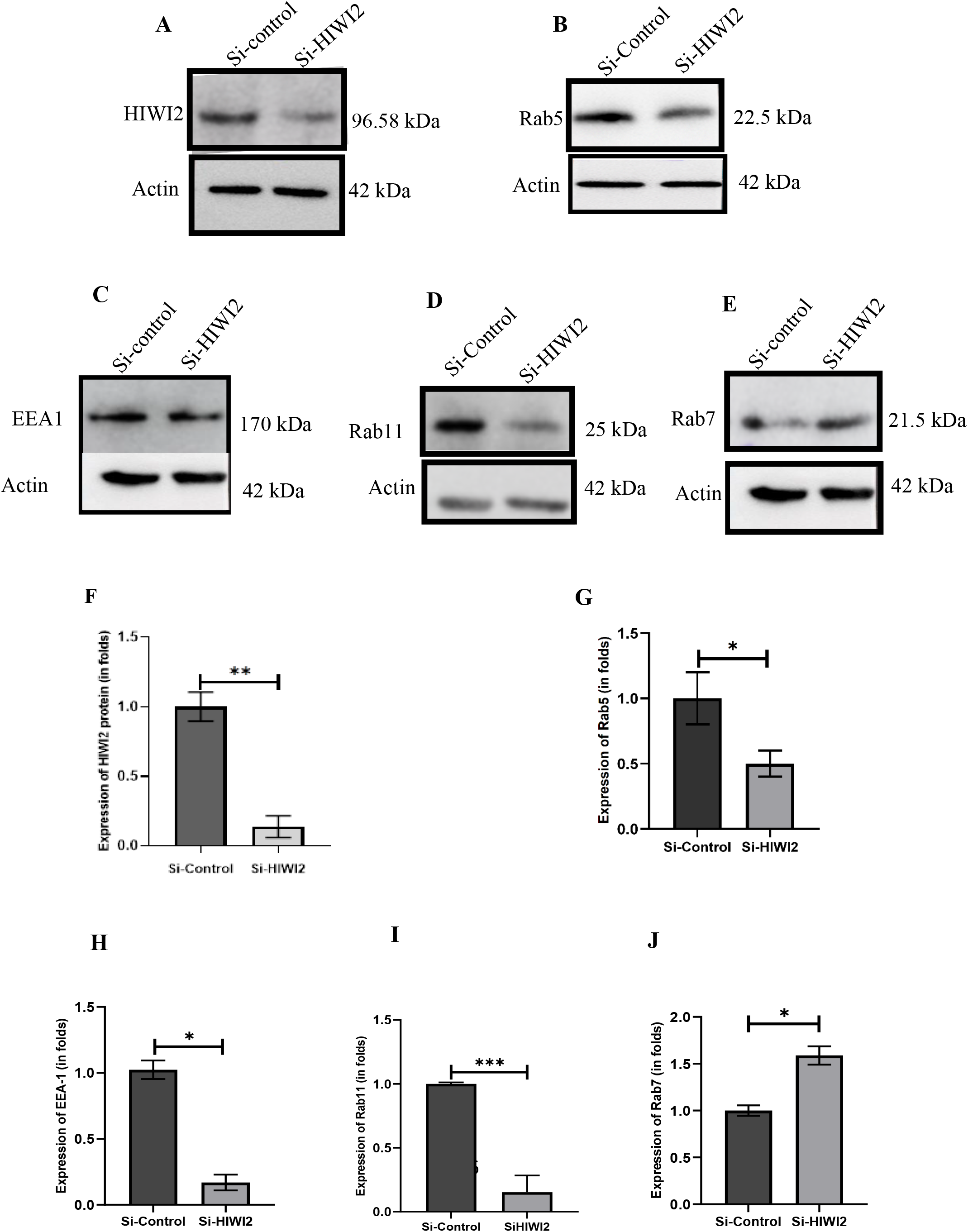
HIWI2 affects the early, recycling and late endosomal proteins associated with trafficking. Knockdown of HIWI2 in 661W cells was checked by Western blot and quantified (A,F). Western blot analysis and quantification of early endosomal protein Rab5 (B,G), early endosomal antigen 1 (EEA1) (C, H), recycling endosomal protein Rab11 (D,I), late endosomal protein Rab7 (E,J). Graph pad prism software was used for quantification. The students’ t-test was used for statistical analysis. Values are means ± SEM, n = 3. *p < 0.05 and **p < 0.01, ***p < 0.001 were considered statistically significant. Original blots for Fig.1A, B, C, D and E are given in supplementary (Fig.S1A, B, C and D).

### 2. HIWI2 knockdown affects an array of Receptor Tyrosine kinases, with Eph receptors most affected

A Phospho RTK-array (ARY001B, CST) was employed to comprehensively assess the impact of HIWI2 on phospho RTK expression in PR. As observed in (Fig.2A) significant altered phosphorylation was observed in various RTKs like Axl (B21,22) Dtk, (B23,24), IGFR(B19,20), HGFR (C5,C6), PDGFR (C9,C10), Tie1(C23,C24), Tie1/2 (D1,D2), NGFR (D5,D6) and most strikingly the Eph family (panel E and F7, F8) had shown significant altered phosphorylation. Specifically, Eph A2, EphA4, EphB2, EphB4 receptors showed substantial decrease in phosphorylation, indicating strong sensitivity to HIWI2 loss. To further validate the data, the expression of EphA2 and EphB2 receptors, which are crucial for the retina and photoreceptors, was assessed. The expression of EphA2 was significantly reduced following the silencing of HIWI2, as observed in Fig.2B. The total expression of EphA2 was reduced by 2-fold as compared to the control. The phosphorylated form of EphA2 was more significantly reduced by 7.2-fold. Similar results were observed for EphB2, as the total and phosphorylated forms of EphB2 were reduced following the silencing of HIWI2 by 2-fold and 2.3-fold, respectively.

**Fig. 2.**
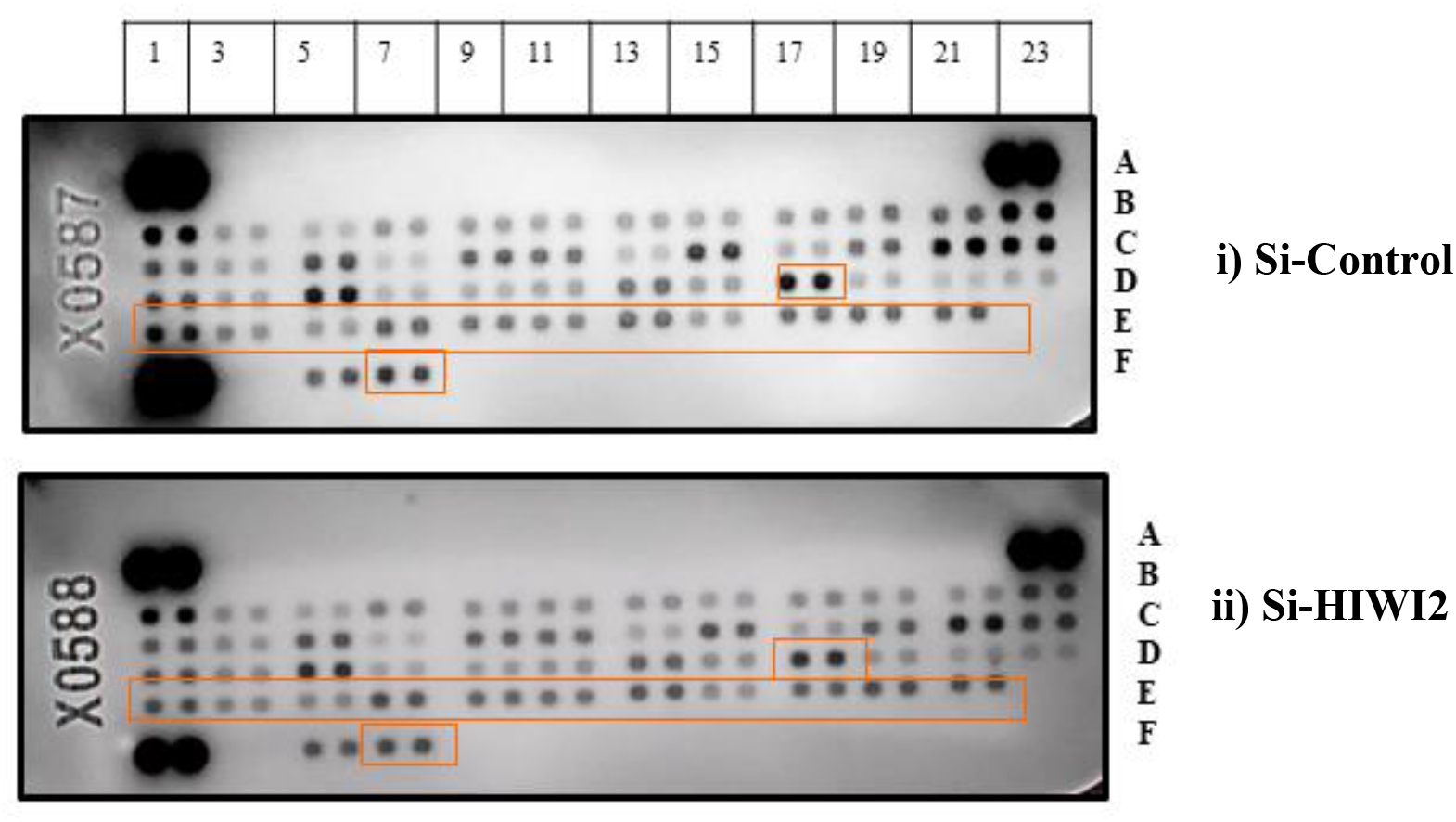
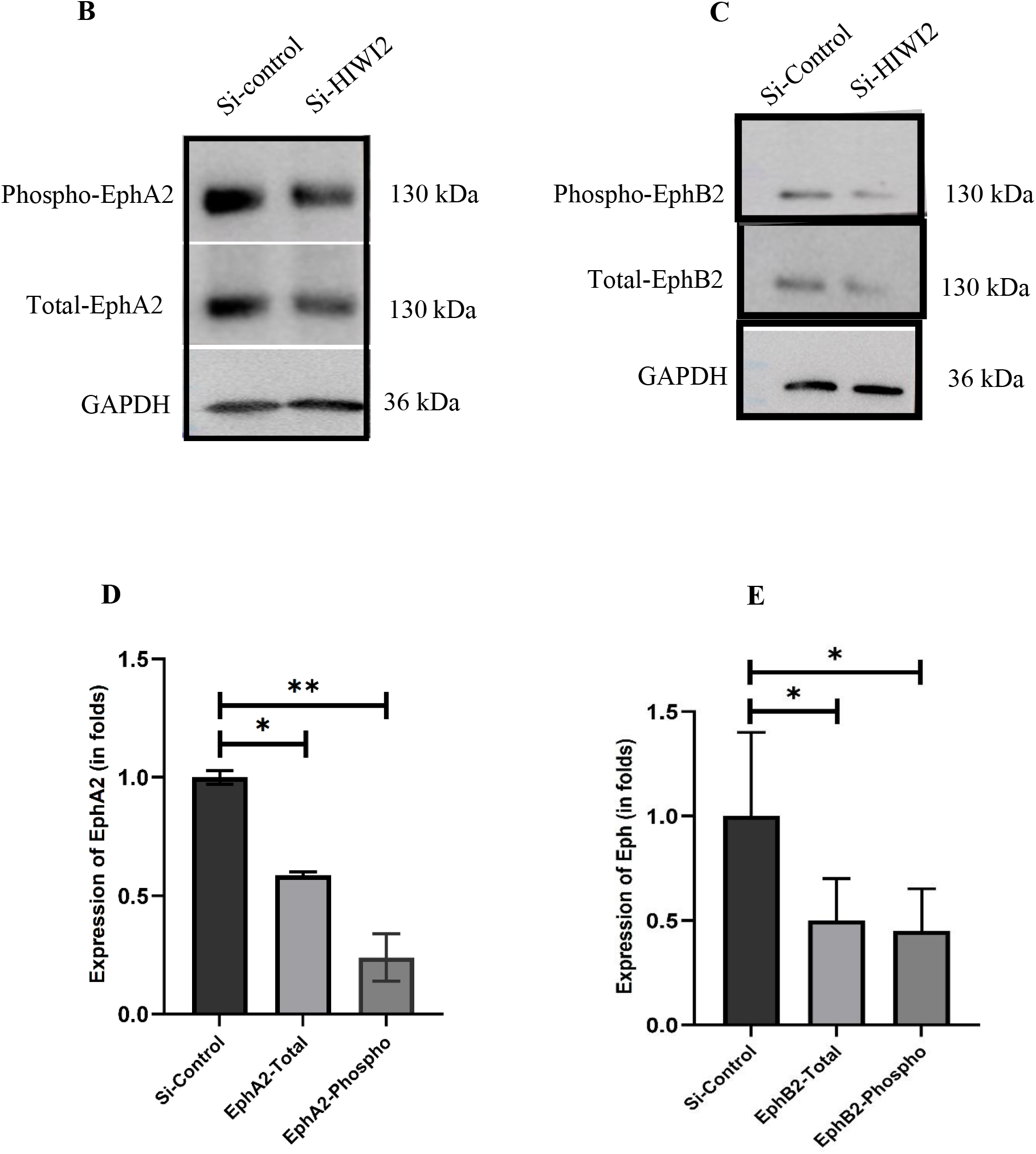
Silencing of HIWI2 alters a range of Receptor Tyrosine Kinases (RTKs). **A)** Si-Control and Si-HIWI2 661W cell lysates were added to Phospho-RTK arrays. Spots are in duplicate and each pair corresponds to a specific Phospho-RTK. **i)** The top panel corresponds to control, and **ii)** while the bottom one corresponds to Si-HIWI2 cells. Phospho-Ephs corresponds to the doublet at E and F rows (marked with red box) showing decreased expression after knockdown of HIWI2. **B)** and **C)** Western blot analysis and **D)** and **E)** quantification of total and phospho form of EphA2 and EphB2 respectively upon knockdown of HIWI2 in 661W cells. The results were quantified and generated by Graph pad prism software. The students’ t-test was used for statistical analysis. Values are means ± SEM, n = 3. *p < 0.05 and **p < 0.01, ***p < 0.001 were considered statistically significant. Original blots for Fig.2B, and C, are given in supplementary Fig.S2B, and C.

### 3. Silencing of HIWI2 results in an elevated expression of Akt as a compensatory response to defective Eph signaling

The Akt and Eph are one of the major regulators of cellular homeostasis, integrating vesicular transport, protein synthesis, and metabolism. These highly interconnected pathways mediate robust survival and stress adaptation signals that are essential for maintaining cells homeostasis. Given the known feedback between Eph signaling and the Akt pathway, we examined Akt activation. HIWI2 depletion induced a 1.68-fold increase in phospho-Akt (p < 0.01), with total Akt unchanged (Fig. 3A,B). This suggests compensatory pro-survival signaling in response to diminished RTK/Eph activation.

**Fig. 3.**
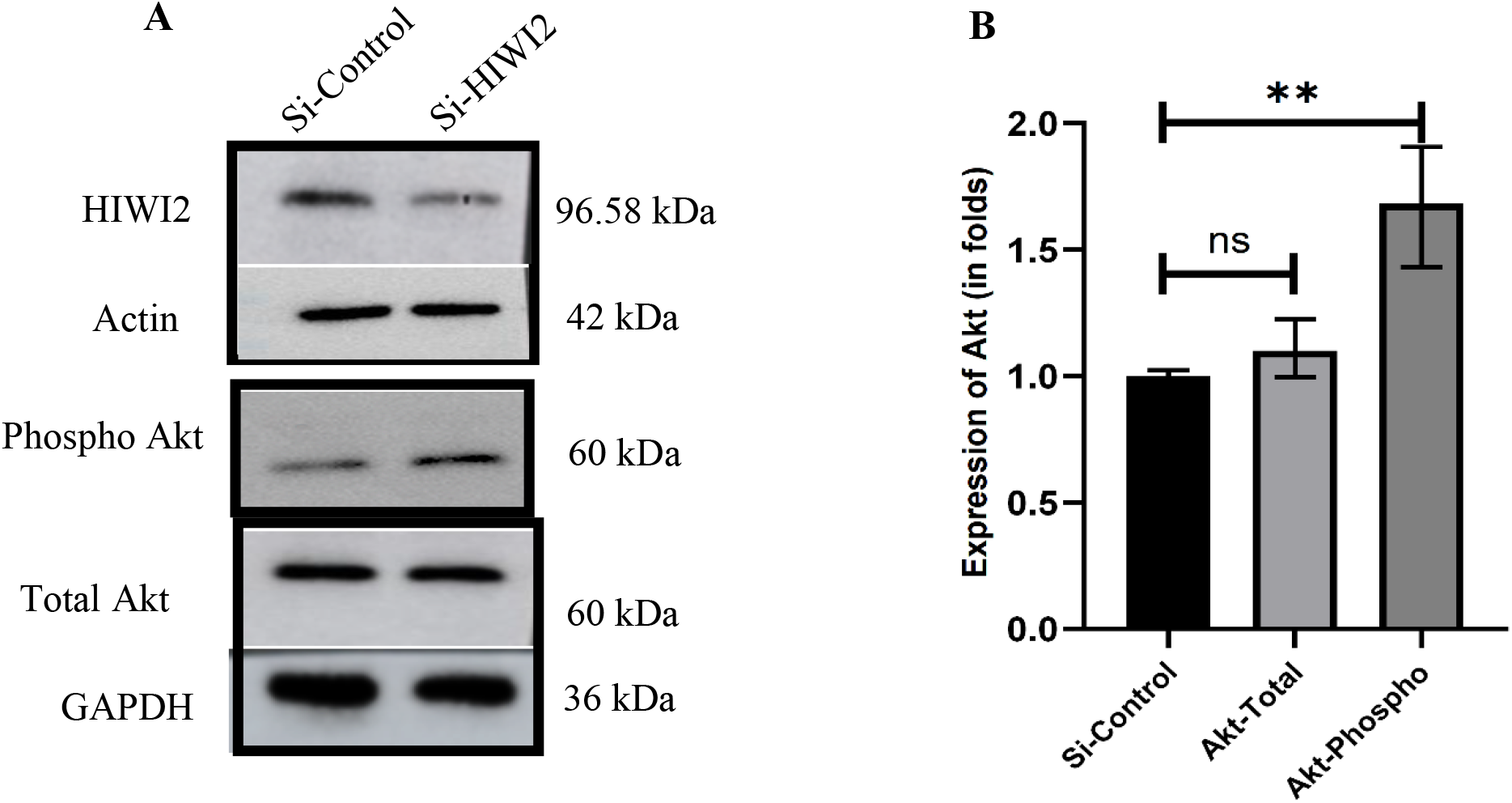
HIWI2 alters the downstream effector molecule Akt. **A) and B)** Western blot analysis and quantification of total and phospho form of Akt. Si-HIWI2 cells showed a significant increase of Phospho-Akt. The students’ t-test was used for statistical analysis. Values are means ± SEM, n = 3. *p < 0.05 and **p < 0.01, ***p < 0.001 were considered statistically significant. ns = not significant. Original blots for Fig.3A are given in supplementary Fig.S3A.

### 4. HIWI2 depletion delays wound closure in photoreceptor cells

We performed an in vitro scratch assay in photoreceptor cells to assess the effect of the alterations in endosomal trafficking, RTK recycling and Akt signaling. HIWI2-silenced PR, displayed a 45% delayed wound closure (p < 0.001) (Fig. 4A), indicating reduced motility-associated behaviour. Although PR are not migratory in vivo, this experiment provides a useful readout of cytoskeleton and membrane trafficking dependent cellular dynamics. These findings suggest that HIWI2 may influence Akt-mediated other downstream pathways that are essential for photoreceptor cell function.

**Fig. 4.**
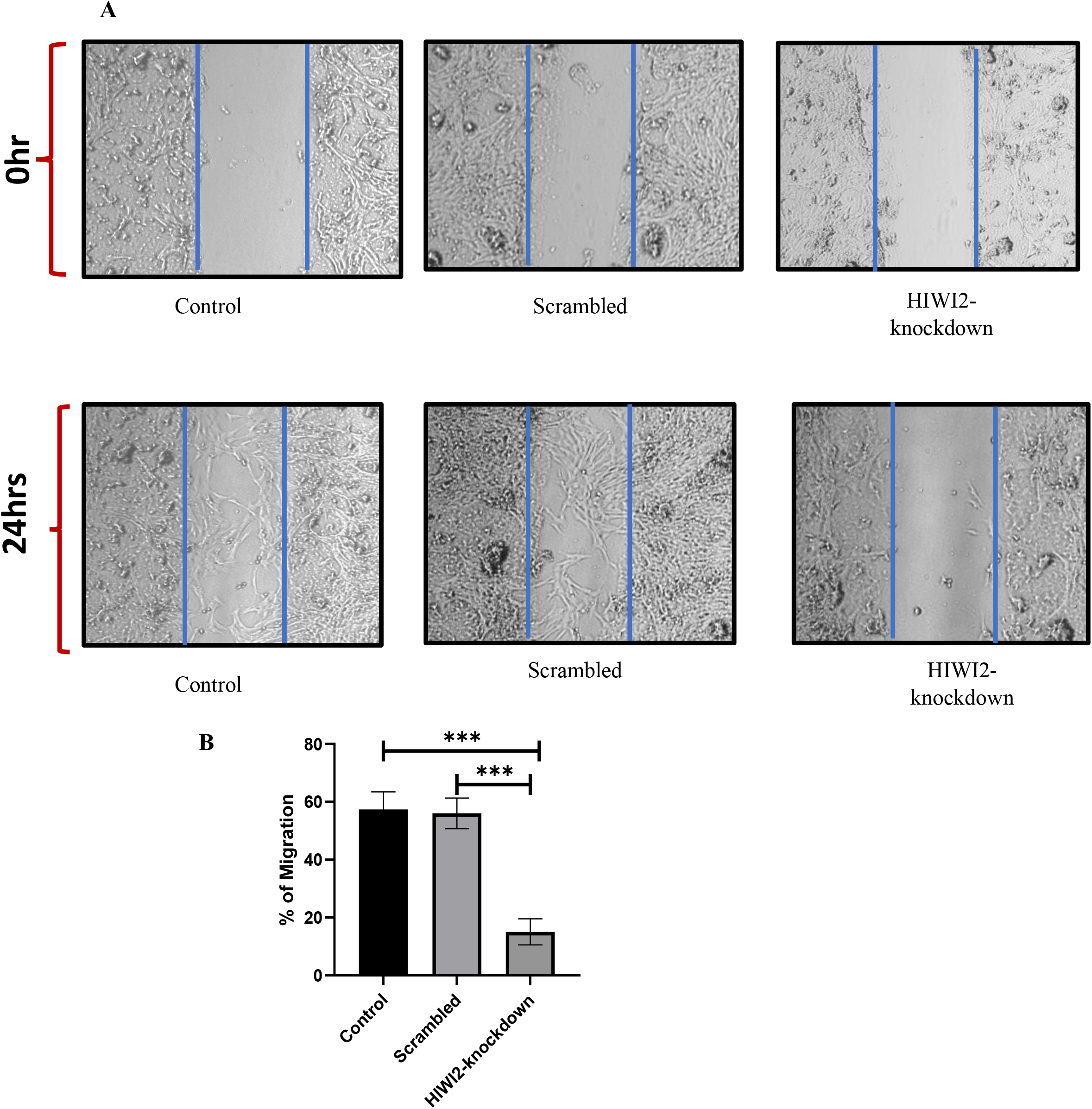
HIWI2 knockdown impairs wound closure in photoreceptor cells. **A)** Scratch was made in Control, Scrambled and Si-HIWI2 661W cells and phase-contrast images were taken at 0 hr and 24hrs, **B)** The bar graph represents the significant delay in wound healing after HIWI2 knockdown in PR. The students’ t-test was used for statistical analysis. Values are means ± SEM, n = 6. *p < 0.05 and **p < 0.01, ***p < 0.001 were considered statistically significant (4x magnification; scale bar, 200 µm).

**Fig. 5.**
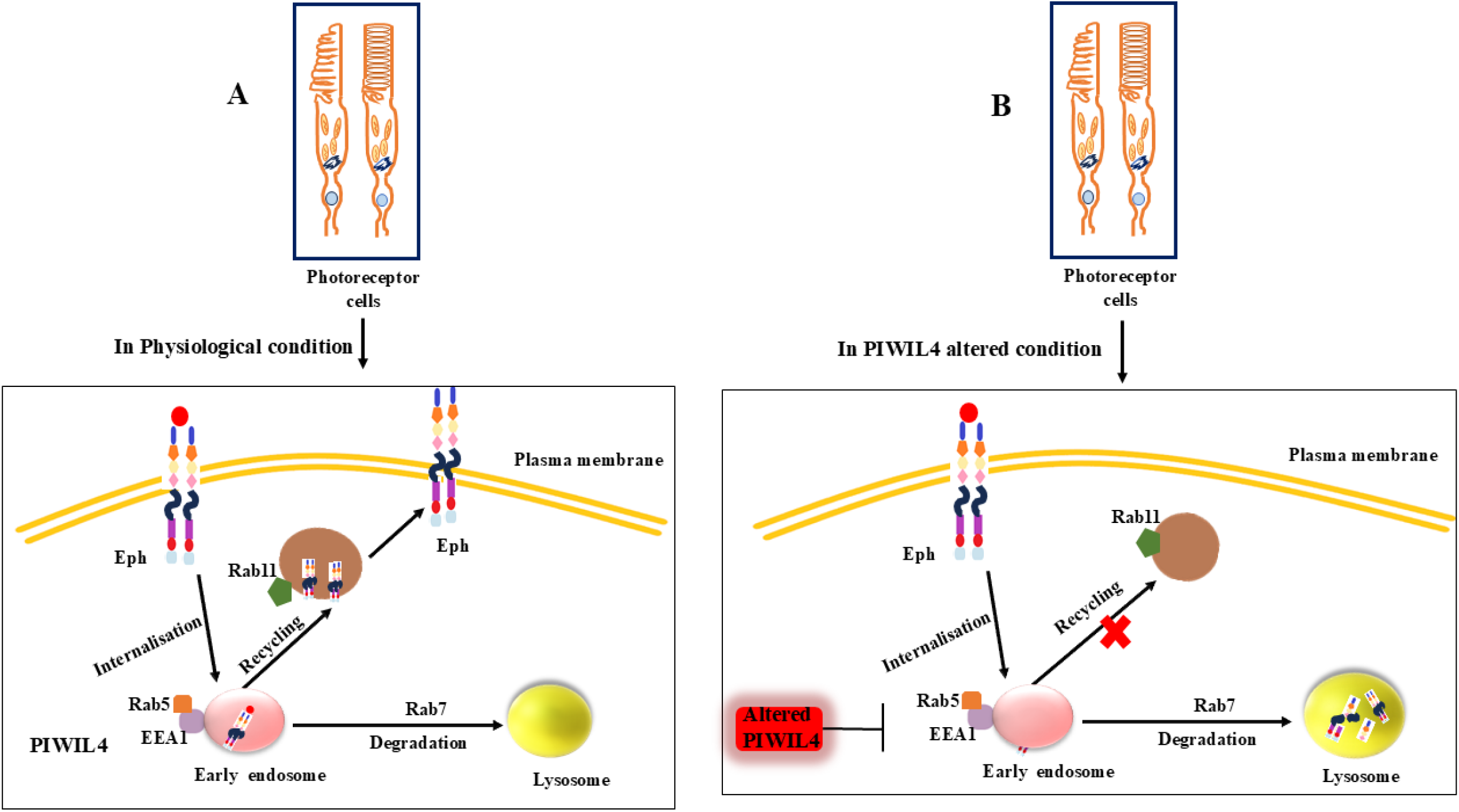
A mechanistic overview of HIWI2-Rab-RTK regulatory network. This figure outlines the mechanistic interplay between the PIWI-like protein HIWI2, the Rab family of small GTPases (Rab5, Rab7, Rab11), and the Eph family of Receptor Tyrosine Kinase. **A) Physiological State**: In normal conditions, HIWI2 represents a crucial modulator within the endosomal network, maintaining appropriate expression/activity levels of Rab5 (early endosomes), Rab7 (late endosomes/lysosomes), and Rab11 (recycling endosomes). Proper balance among these Rabs promotes optimum trafficking of internalized Eph receptors, allowing the proper proportion of Eph signaling receptors to be recycled back to the plasma membrane versus those subjected to degradation via lysosomes, to maintain signaling homeostasis. **B) Perturbed State:** In contrast, experimental suppression or functional elimination of HIWI2 disrupts this regulatory function, with loss of expression of specific Rab GTPases shown here. This quantitative Rab deficiency leads to compromised sorting fidelity within the endosomal network, substantial disruption in the rates of Eph receptor trafficking and recycling, leads to an altered Eph signaling homeostasis and the emergence of dysregulated cellular phenotypes.

## Discussion

This study reveals HIWI2 as a previously unidentified regulator of PR membrane trafficking and receptor homeostasis. HIWI2 knockdown results in a significant downregulation of multiple receptor tyrosine kinases (RTKs), including Eph, VEGFR, EGFR, Axl, etc, impacting a broad range of cellular signaling pathways and functions.

Previous studies have established that Rabs are essential in vesicle trafficking as well as membrane fusion processes. Deretic et al. [37] were the first to establish the function of Rabs in directing rhodopsin to the PR cilium. This was later clarified by Wang and Deretic [38], who established the Rab11-Rab8 cascade in protein trafficking to the specialized ciliary outer segment. These findings clearly indicate the importance of Rab-mediated transport and ciliary function in the renewal of the outer segment, thereby affecting PR function, structure, and the prevention of retinal degeneration [39].

The established involvement of Rabs in ocular development, their pivotal regulatory role in RTK signaling [40–42], and the observation from this study of HIWI2-mediated RTK modulation, indicates the impact of HIWI2 silencing on Rabs expression in PR. Interestingly here we found that HIWI2 loss caused a coordinated reduction in early and recycling endosomal regulators (Rab5, EEA1, Rab11) and an increase in Rab7, shifting trafficking away from processing/recycling and toward degradation. This trafficking imbalance resembles known degeneration pathways in PR where Rab5 and Rab11 deficiencies misroute rhodopsin and disrupt outer segment maintenance [19,43], whereas Rab7 elevation accelerates degradative flux and contributes to advanced degeneration [44–45].

The inherent spatiotemporal precision of RTK signaling prompted an investigation into the role of ncRNAs and RNA-binding proteins (RBPs) in regulating RTKs expression. Existing literature suggests that ncRNAs and RBPs can modulate RTK expression and signaling in response to cellular stress [46,47]. A central finding in this study is the strong decrease in Eph receptor activation following HIWI2 depletion. Eph receptors are significant for retinal development, lamination, and synaptic targeting [48,49]. Because RTK signaling also depends on Rab-mediated trafficking [50], the altered Rab landscape provides a mechanistic explanation for EphA2/B2 dysfunction. This possibly reduces the receptor stability and phosphorylation, enhancing the lysosomal routing and impaired recycling, consistent with the loss of Rab11 and an elevated Rab7.

In this study, we have focused on the steady-state protein levels to determine the effect of HIWI2 knockdown. Our biochemical analysis by Western blotting has clearly shown the change in Eph receptor and Rab protein expression after HIWI2 knockdown, however experiments using high-resolution immunofluorescence imaging would be helpful to supplement our results by precisely locating the trafficking vesicles where these interactions take place.

The serine/threonine kinase Akt plays a pivotal role in cellular regulation, modulating cell size, proliferation, and survival via downstream effectors, notably mTORC1. Canonical RTK signaling activates Akt through PI3 kinase, initiating a cascade that culminates in Akt phosphorylation, thereby activating it [51]. Conversely, Eph receptor forward signaling has been reported to suppress Akt activation [52–54]. Notably, Akt phosphorylation was found to be increased after HIWI2 knockdown in PR cells. Similar increases in Akt activation in retinal pigment epithelial cells were also observed following HIWI2 depletion [55] and tight junction disruption, suggesting that enhanced Akt expression may be a common retinal response to HIWI2 loss and indicating that HIWI2 could play a role in maintaining broader retinal cellular equilibrium. Activation of Akt, typically linked to pro-survival signalling, did not translate into improved cellular behaviour in this scenario, implying it may be a compensatory response to disrupted receptor stability and a trafficking imbalance rather than increased functional integrity [56–58].

PR motility is defined not by cellular migration, but by structural dynamics. This constant remodeling and intracellular transport are fundamental to light adaptation, photon capture efficiency, and the overall architectural stability of the retina [59]. After demonstrating that HIIWI2 silencing disrupts these dynamics, the next step would be to elucidate the underlying signaling architecture that sustains PR motility and its structural integrity. It would be interesting to further investigate proteomic profiling that could help map the specific effector proteins regulated by HIWI2 in the PR cytoskeleton.

Overall, our research reveals a novel regulatory mechanism wherein HIWI2 maintains Rab-dependent endosomal homeostasis to ensure proper Eph receptor activation and downstream Akt-mediated compensatory pathway. Collectively our study suggests that HIWI2 as a potential player in the intricate balance of RTK signaling and cellular trafficking within PR.

## Conclusion

This research identifies a previously unknown functional synergy between HIWI2 and Rab proteins within PR. By demonstrating how HIWI2 bridges the gap between RNA-binding mechanisms and vesicular trafficking, the study underscores its essential role in maintaining the PR integrity. The functional interaction of HIWI2, Eph, and Akt signaling indicates a complex regulatory mechanism affecting cellular activities such as cell-cell contact, and different stress-related pathways. It also suggests that HIWI2 is essential for maintaining the delicate balance of signaling pathways involved in effective PR function and retinal health. Future studies using in vivo models, neuronal differentiation, and synaptic marker analyses will further elucidate HIWI2’s role in retinal physiology and degeneration.

## Acknowledgement

We acknowledge the support provided by the Science and Engineering Research Board Extramural Grant (SERB-EMR) file no. EMR/2017/000002 and Indian Council of Medical Research (ICMR) project NO. 5/4/6/1/OPH/2015-NCD-II. The funding body is not involved in the design of the study and collection, analysis, and interpretation of data and in writing the manuscript. RR thanks the Department of Science and Technology (DST) for the fellowship under project NO.SR/WOS-A/LS-388/2018. SC thanks the support from the University Grants Commission, New Delhi for the award of Assistant Professorship under its Faculty Recharge Program (UGC-FRP).

## Author Contribution

**Subbulakshmi Chidambaram:** Writing – Review & editing, Supervision, Resources, Project administration, Funding acquisition, Conceptualization. **Rupa Roy:** Writing – Original draft, Experiments, Data curation, Resources, Conceptualization. **Jayamuruga Pandian Arunachalam** – Review with critical inputs, Resources. **Rahini Rajendran** – Resources. All authors read and approved the final manuscript.

## Declarations

### Funding

Science and Engineering Research Board Extramural Grant (SERB-EMR) file no. EMR/2017/000002.

Indian Council of Medical Research (ICMR) project NO. 5/4/6/1/OPH/2015-NCD-II.

University Grants Commission, Faculty Recharge Program (UGC-FRP). New Delhi.

Department of Science and Technology-Women Scientist Fellowship (DST); project NO.SR/WOS-A/LS-388/2018

### Declaration of generative AI and AI-assisted technologies in the writing process

During the preparation of the manuscript, AI tools were used only for limited purposes such as to improve the language and readability or restructuring sentences. After using the tool, we reviewed and edited the content as needed and took full responsibility for the publication’s content.

### Ethics Approval

Not applicable

### Informed Consent

Not applicable

### Data availability

All data supporting the findings of this study are available within the paper and its Supplementary Information.

